# The *Streptococcus pyogenes* M protein is involved in phenotypic resistance to phage A25 infection in presence of human serum

**DOI:** 10.1101/2024.02.07.579182

**Authors:** Lionel Schiavolin, Jenny Steinmetz, Gwenaëlle Botquin, Valérie Delforge, Dalila Lakhloufi, Pierre R. Smeesters, Anne Botteaux

## Abstract

*Streptococcus pyogenes* is responsible for mild to life-threatening infections. Bacteriophages, or phages, and their virulence genes play a key role in the emergence and expansion of epidemics. However, relatively little is known about the biology of *S. pyogenes* phages, particularly in biologically relevant environments. During infection, *S. pyogenes* conceals from the host immune system through the binding of human serum proteins. This evasion is mediated by surface proteins, such as the M protein which is a major virulence determinant of *S. pyogenes.* Here, we demonstrate that human serum proteins also confer phenotypic resistance to phage A25 infection by impeding phage adsorption. We have found that, although not directly involved in phage A25 infection, the M protein is involved in this inhibition through the binding of both IgG and albumin, especially in absence of bound fatty acids. These findings highlight the importance of studying phages within a physiological context, specifically in the environmental conditions in which they will be used.

**Author summary:** The issues of antimicrobial resistance and resurgence of life-threatening infection, like the recent cases of invasive *S. pyogenes* infections, are prompting the scientific community to use phages as a complementary therapy. Phages are often characterized in laboratory conditions which are very different from the infection site. During human infection, *Streptococcus pyogenes* uses serum proteins to protect against the immune system. Our data illustrate how the human host environment also modulates phage susceptibility of *S. pyogenes*. We found that human serum transiently protects a M25 strain against infection by the lytic phage A25. This protective effect is mediated in part by the M protein, a major virulence determinant and the target of current vaccines. This new function for the M protein highlights the need to characterize bacteria-phage interactions in a more physiological context to increase the chances of success of phage therapy.

## Introduction

*Streptococcus pyogenes*, or Group A *Streptococcus*, is a chain-forming Gram-positive bacterium and obligate human pathogen. *S. pyogenes* asymptomatically colonizes the throat and skin but also causes a wide variety of diseases. These infections range from mild and superficial (pharyngitis, impetigo…) to life-threatening and severe, either toxin-mediated (scarlet fever, toxic shock syndrome…) or invasive (necrotizing fasciitis, cellulitis…) (1). *S. pyogenes* is responsible for more than 727 million superficial infections, 1.78 million severe infections and at least 500,000 deaths yearly (2). Infections are treated exclusively with antibiotics. Although *S. pyogenes* has so far remained susceptible to penicillin, treatment failures have been reported (3). Additionally, resistance to macrolides and tetracyclines is common in several geographic regions (4). Treatment failure, antimicrobials resistance and high mortality associated to severe infections highlight the need for alternative treatment options until a vaccine is available. Amongst the alternatives, bacteriophages and their products have regained interest during the last decades (5–7).

Bacteriophages (phages in short) are viral predators of bacteria. Based on their life cycle, they can be divided into temperate and virulent phages. For both types, infection starts with phage adsorption to their host, through the binding of bacterial receptors, and injection of their genetic material. According to environmental and host metabolic cues, temperate phages can either enter in a ‘dormant’ state (lysogeny) or in a lytic cycle characterized by host hijacking, virions production, and release following bacterial lysis. In contrast, virulent phages can only perform the lytic cycle. During lysogeny, phage can insert their DNA in the host genome (prophage) which may provide a selective advantage to the newly formed lysogen (8). Bacteria encode a plethora of immune mechanisms to defend against phage infection, the most described being the Restriction-Modification and CRISPR-Cas systems which targets the injected phage genome. Another way to avoid phage infection is to prevent their adsorption by mutating, modifying, masking receptors or by using decoys (9).

For *S. pyogenes*, few virulent phages have been isolated and characterized. Amongst them, the phages A1 and A25 have been the subject of recent studies (10–12). The phage A25 (φA25) is a highly transducing phage with an average burst size (12-30 plaque forming units (PFU)/cell) that varies according to the host strain and growth medium used (10,13). Genomic and functional studies in *S. pyogenes* have shown that temperate phages carry a plethora of virulence genes which confer an increased resistance to the human immune system (10,14,15). Moreover, population genomics have highlighted their key role in the expansion of different pandemic lineages in the eighties (16). Although the anti-phage CRISPR-Cas9 system has been discovered in *S. pyogenes*, little is known about the biology of its phages and even less in the context of the human host (10–12,17,18).

During infection, *S. pyogenes* encounters the human immune system. Notably, an increase in the local concentration of plasma proteins occurs during tonsillitis (19). *S. pyogenes* is weaponized with surface proteins that possess the capacity to bind plasma proteins (20). These interactions form a shield-like structure at the bacterial surface, preventing the deposition of opsonins and protecting against antimicrobial peptides (20,21). The surface M protein is the main protein involved in this process, and represents the principal target of vaccine development as well as the basis for *S. pyogenes emm*-typing (22). During growth in human plasma or serum, the M1 protein is able to bind to human serum albumin (HSA), allowing the acquisition of exogenous fatty acids (23). Based on a functional classification of M proteins (24), the M25 protein (*emm*-cluster E3) is predicted to non-specifically bind IgG. Considering the role played by plasma/serum during *S. pyogenes* infection and the formation of a shield-like structure at its surface, there is a need to address the biology of *S. pyogenes* phages in this context which will inform about their potential use in therapy.

Here, we investigated the infection of a *S. pyogenes* M25 strain by the virulent φA25 *in vitro* and *ex vivo*, in presence of human serum. We found that the presence of human serum reduces the adsorption of φA25 on *S. pyogenes* preventing its dissemination in the bacterial population. We identified the M protein as taking part in this process. The binding of immunoglobulins or of fatty acid-free HSA partially blocked φA25 infection while we found that a combination of both serum proteins reduces infection at a significant level. Collectively, our data illustrate the requirements to study and characterize phages in a more bench-to-bedside way.

## Results

### Determination of φA25 infection parameters

To study the infection by the phage A25 (φA25), we first determined its infection parameters on a susceptible *S. pyogenes* M25 strain (NCTC8306) in Todd-Hewitt (TH) medium supplemented with 0.5% Yeast extract (THY) at 37°C. The φA25 adsorbed in a two-step process characterized by a rapid adsorption of 60% of the phages population within the first two minutes and a second slow-rate adsorption. Ultimately, more than 80% of phages adsorbed within the first ten minutes (Fig 1A). A one-step growth experiment was then performed to determine the timing of phages production and release. We detected functional phages 31min post-infection (eclipse) and they started to be released after 36min (latency) with a burst size of approximately 25.8±3.2 (Fig 1B). We next determined the multiplicity of infection (MOI), *i.e.* the number of phages per bacteria, required to eradicate the whole population. As *S. pyogenes* is a chain-forming bacterium, we first sonicated bacteria to dissociate chains (Fig S1A). Isolated cells were then infected for 10min with increasing MOIs of phages and plated on THY agar for CFU counting. We found that a MOI of 25 is sufficient to kill most bacteria (Fig S1B). We then infected exponentially grown bacteria with a MOI of 25 and monitored infection in real-time by measuring optical density (OD) at 600nm. We found that the population was lysed in one lytic cycle (Fig 1C). Finally, we performed a growth/lysis assay by co-culturing the bacterial population with phages at different MOIs and measured OD every 10 minutes. We found that a MOI of 0.001 was sufficient to collapse the bacterial population in less than 5 hours (Fig 1D).

**Fig 1.**
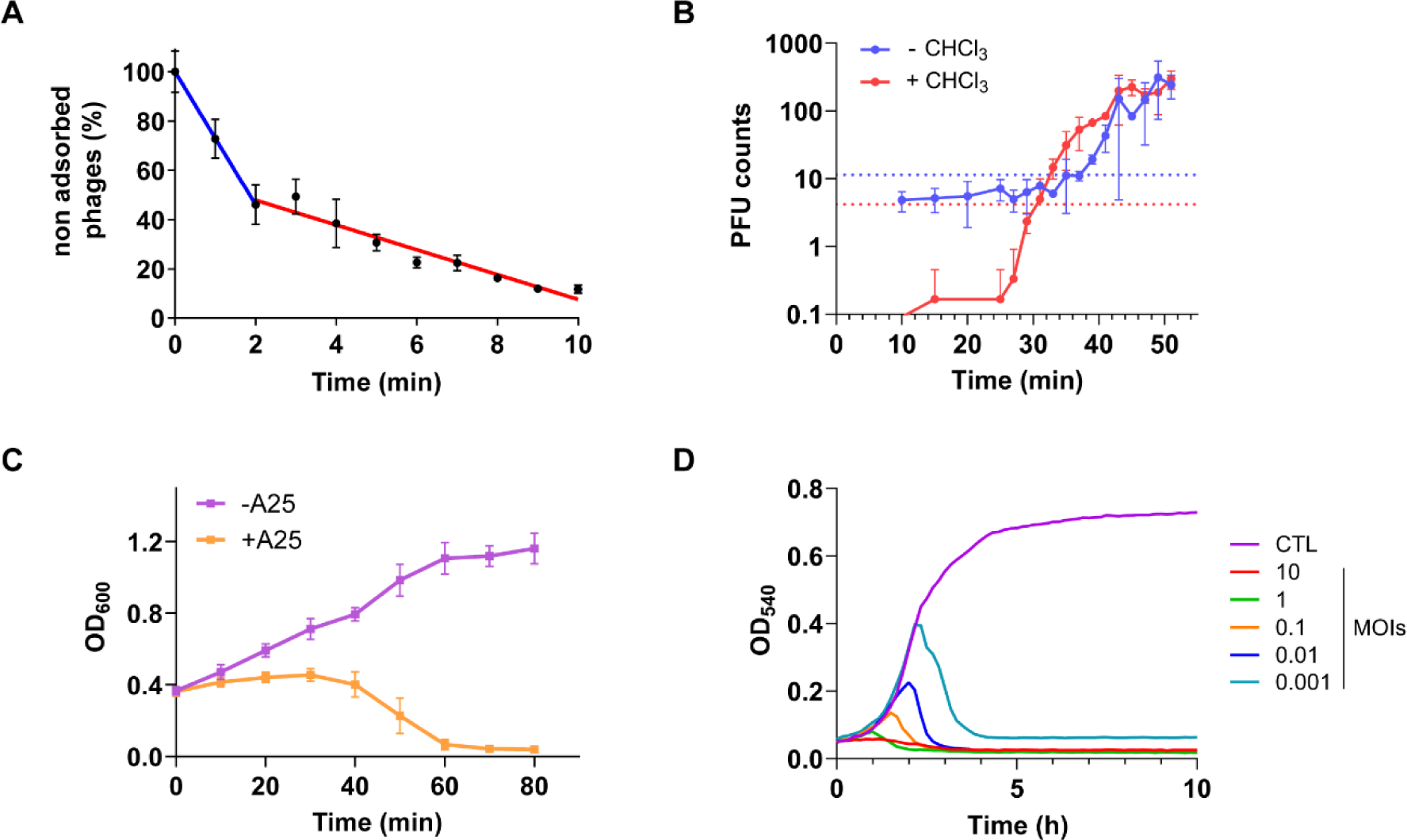
Phage φA25 – M25 strain infection parameters. **(A)** φA25 adsorption on the *S. pyogenes* M25 strain in THY medium (MOI=0.001). Linear regressions were applied to experimental data (0 to 2min, *blue* and 2 to 10min, *red*). **(B)** One-step growth curve of φA25 in the *S. pyogenes* M25 strain grown in THY medium (MOI=0.01). Free phages (+CHCl_3_, *red*) and free phages + infective centres (-CHCl_3_, *blue*) were quantified by PFU counting. Dotted lines correspond to 5% of total PFU counts. Times where dotted and solid lines cross correspond to eclipse (+) and latency (-) periods. **(C)** Growth (purple) and collapse (orange) curves with the φA25 (MOI=25) of *S. pyogenes* M25 previously grown to exponential phage in THY medium. **(D)** Growth and collapse curves with φA25 at different MOIs (as indicated) of *S. pyogenes* M25 in TH medium. Data (A-C) are displayed as mean ± SD of at least three independent experiments. Data in D are a representative of three independent experiments displayed in Fig S1C.

### φA25 infection is prevented by human serum

During tonsillitis, *S. pyogenes* faces an increase in the local concentration of plasma/serum proteins (19). Since *S. pyogenes* is able to bind these proteins to hide from the host immune system (20,21), we reasoned that they could also mask phage receptors. We therefore infected bacteria in TH medium supplemented with 20% serum, which ranges to plasma concentrations found during tonsil infection (19). We found that the addition of human serum dramatically reduced bacterial collapse, even for the highest MOI that prevents bacterial growth in TH medium (Fig 2A, to compare with Fig 1D).

**Fig 2.**
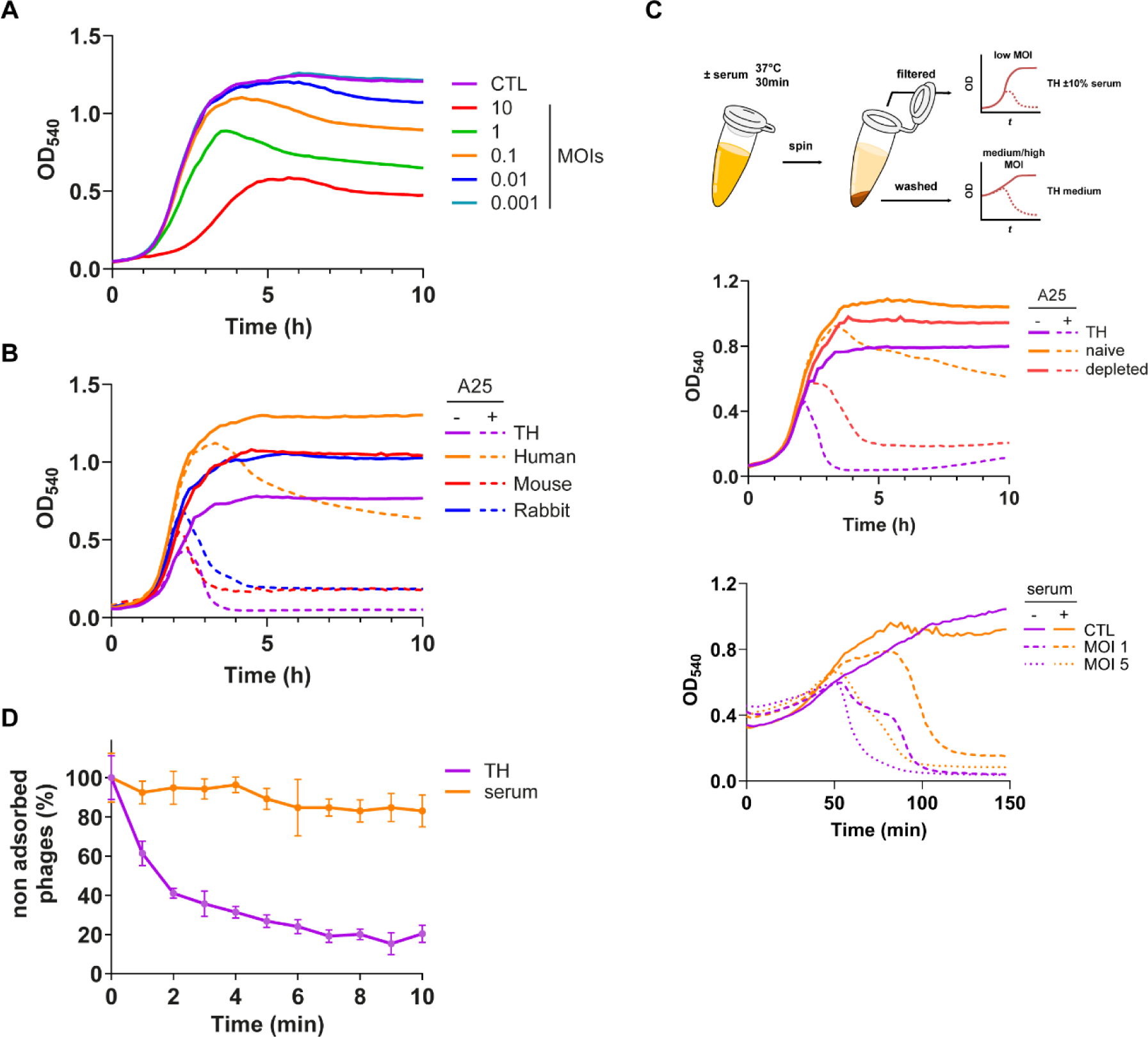
Human Serum protects against φA25 infection. **(A)** Growth and collapse curves with φA25 at different MOIs (as indicated) of *S. pyogenes* M25 in TH medium supplemented with 20% serum. **(B)** Growth (*solid line*) and collapse (*dotted line*) curves with φA25 (MOI=0.01) of *S. pyogenes* M25 in TH medium alone (*purple*) or supplemented with 20% Human (*orange*), Mouse (*red*) or Rabbit (*blue*) serum. **(C)** Schematic representation of the experiment where exponentially grown bacteria are preincubated with TH medium alone or 100% human serum (*upper panel*). *Central panel*, growth (*solid line*) and collapse (*dotted line*) curves with φA25 (MOI=0.01) of *S. pyogenes* M25 wild-type (WT) in TH medium supplemented or not (*purple*) with 10% naive (*orange*) or depleted (*red*) serum from the preincubation step. *Lower panel*, untreated (*purple*) and serum-treated (*orange*) bacteria from the preincubation step were washed and grown (*solid line*) or infected (*dotted lines*) in TH medium at low to medium MOI as indicated. **(D)** φA25 adsorption on the *S. pyogenes* M25 strain (MOI=0.001) in TH medium alone (*purple*) or supplemented with 20% human serum (*orange*). Data (A-C) are a representative of three independent experiments displayed in Fig S2A-C. Data in D are displayed as mean ± SD of three independent experiments.

*S. pyogenes* is an obligate human pathogen, yet several animal models have been successfully used to study its pathogenesis (25). To determine if the protective effect of serum can be reproduced in animal models, we further used mouse and rabbit sera at the same concentration. We found that both sera were unable to protect the bacterial population against φA25 infection (Fig 2B). To determine whether protection occurs on the bacterial side, exponentially grown bacteria were pre-incubated with serum, washed and infected at different MOIs. We observed a lysis delay for serum-treated bacteria (Fig 2C, *lower panel*). Moreover, we performed a protection assay using the spent serum obtained from this experiment. We observed a reduced protection compared to “naive” serum (Fig 2C, *middle panel*), suggesting that serum components involved in protection were depleted during the preincubation step. Finally, to determine which step of the infection is impaired, we performed an adsorption experiment by concomitantly adding serum and phages to bacteria. We found that the amount of non-adsorbed phages remained high (83.0±8.2%) in presence of serum compared to CTL condition (20.4±4.4%) (Fig 2D). The immediate observation of protection upon serum addition suggests that serum conceals the bacterial receptor rather than reducing its surface expression. These data therefore suggest that the binding of human serum proteins to the bacterial surface provides phenotypic resistance through masking φA25 receptor.

### The M protein is involved in the serum-mediated protection against φA25 infection

We next intended to identify which bacterial protein is involved in preventing φA25 adsorption. As a Gram-positive bacterium, *S. pyogenes* possesses a cell wall with a thick peptidoglycan (PG) which serves as an anchor for teichoic acid, the Group A Carbohydrate (GAC) and surface proteins. These proteins are secreted by the Sec system and, in presence of a C-terminal LPXTG-motif, covalently attached to the PG by the sortase SrtA. These LPXTG-containing proteins are involved in adhesion to the host tissue and immune evasion. Amongst them, the M protein coats the entire bacterial surface and several *S. pyogenes* strains also harbour the M-like proteins Enn and Mrp, which all bind several serum proteins (20,23,26). We sequenced the M25 genome and found that the *mga* regulon encodes the *emm25*, *enn83* and *mrp174* genes (unpublished data). We therefore generated a triple KO (TKO) mutant to determine if these major virulence proteins play a role in serum-mediated protection against φA25 infection. We found that the TKO mutant was more susceptible to φA25 infection in presence of serum compared to the WT strain, while no differences were observed in TH medium (Fig S3A). We next generated *emm*, *enn* and *mrp* individual mutants and found that the M protein was the predominant protein responsible for this phenotype as protection was maintained in the *Δenn* and *Δmrp* backgrounds (Fig 3A and Fig S3A-B). Again, no differences were observed in TH medium compared to the WT strain (Fig S3C). We then incubated exponentially grown M25 WT and *Δemm* strains with serum and used the spent sera in a protection assay. We found that the *Δemm* spent serum conferred an intermediate protection against φA25 infection between naive and WT spent sera (Fig 3B). Altogether, these data suggest that the M protein bind serum components which participate in the protection against φA25 infection.

**Fig 3.**
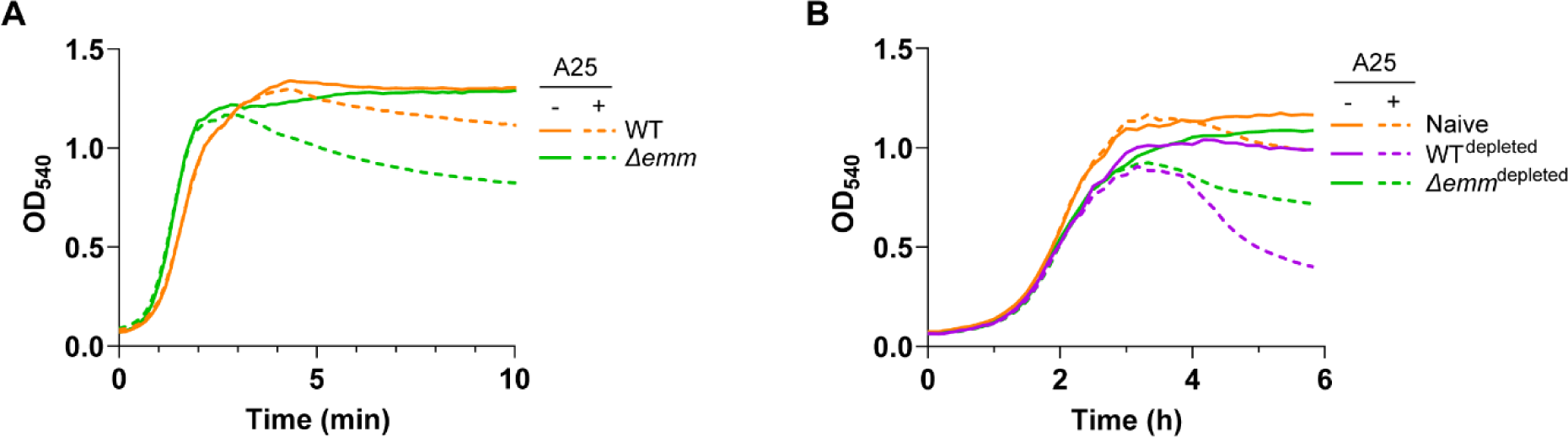
The M protein contributes to serum-mediated protection against φA25 infection. **(A)** Growth (*solid line*) and collapse (*dotted line*) curves with φA25 (MOI=0.01) of *S. pyogenes* M25 wild-type (WT) (*orange*) and *<u>Δemm</u>* deletion mutant (*green*) strains in TH medium supplemented with 20% serum. **(B)** Growth (*solid line*) and collapse (*dotted line*) curves with φA25 (MOI=0.01) of *S. pyogenes* M25 wild-type (WT) in TH medium supplemented with 10% naive serum (*orange*), WT (*purple*) or *<u>Δemm</u>* (*green*) depleted sera were added at a 10% final concentration in TH medium. Data (A-B) are a representative of three independent experiments displayed in Fig S3B and S3D.

### Immunoglobulins and human serum albumin participate in protection against φA25 infection

To further explore the protection mechanism of serum and gain a first insight in the role of serum proteins, we heat-treated serum at 70°C to partially induce proteins unfolding and aggregation, without coagulation. Incubation of bacteria-phages with heat-treated serum reduced protection compared to untreated serum (Fig 4A). We next investigated the binding of serum proteins to the M25 strain using a whole cell binding (WCB) assay and found differences in serum proteins binding (Fig 4B). Notably, we observed a decrease in binding of high molecular weight proteins (>180kDa) and a low molecular protein (>34kDa). To further explore the protection mechanism of serum and gain a first insight in the role of serum proteins, we heat-treated serum at 70°C to partially induce proteins unfolding and aggregation, without coagulation. Incubation of bacteria-phages with heat-treated serum reduced protection compared to untreated serum (Fig 4A). We next investigated the binding of serum proteins to the M25 strain using a whole cell binding (WCB) assay and found differences in serum proteins binding (Fig 4B). Notably, we observed a decrease in binding of high molecular weight proteins (>180kDa) and a low molecular protein (>34kDa). Moreover, we found an increase in proteins that could correspond to human serum albumin (HSA, 69kDa) and immunoglobulins (HC, 50kDa and LC, 25kDa).

**Fig 4.**
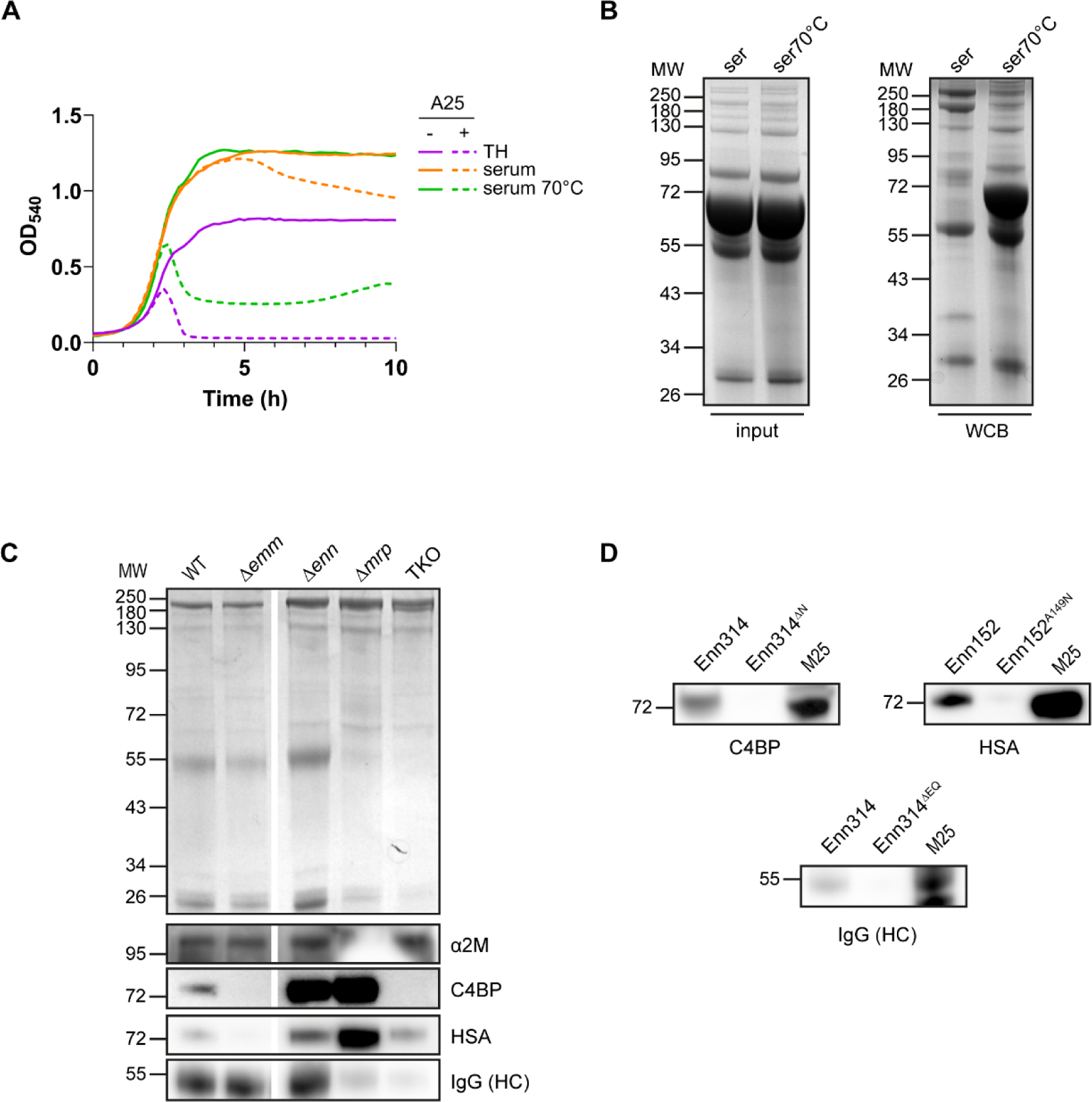
Role of serum proteins in protection against φA25 infection. **(A)** Growth (*solid line*) and collapse (*dotted line*) curves with φA25 (MOI=0.01) of *S. pyogenes* M25 in TH medium alone (*purple*) or supplemented with 20% untreated (*orange*) or heat-treated (*green*) human serum. **(B)** SDS-PAGE/Coomassie blue staining profiles of untreated (ser) or heat-treated (ser70°C) human serum (input, *left*) and of serum proteins bound to the surface of the M25 wild-type (WT) strain (WCB, *right*). **(C)** SDS-PAGE/Coomassie blue staining (*upper panel*) or immunoblotting (*lower panels*) profiles of serum proteins bound to the surface of the M25 wild-type (WT) and mutant (*Δemm*, *Δenn*, *Δmrp* and triple KO) strains. Data are representative of three independent experiments. **(D)** SDS-PAGE/immunoblot of serum proteins eluted with the recombinant M25-His protein from a Ni-NTA resin. Wild-type Enn152, Enn314 and their derivatives were used as controls. MW, *molecular weight* in kilodalton. Data in A are a representative of three independent experiments displayed in Fig S4B. Data (B-D) are representative of three independent experiments. SDS-PAGE/Coomassie blue staining profile of purified proteins (D) is displayed in Fig S4C.

Since the deletion of the M protein decreases serum-mediated protection against φA25 infection, we determined *in silico* binding motifs of the M25 protein based on the literature (24,27–30). We found three HSA- and one IgG-binding motifs (Fig S4A). In addition, the hypervariable region (HVR) of the M25 protein is predicted to bind C4BP (Fig S4A) (30). We also predicted that the Mrp174 is composed of 3 A-repeats known to bind IgG3 (27) and contains three putative IgG-binding motifs (1-3 mismatches), while the Enn83 protein possesses three HSA-binding motifs (0-3 mismatches) (Fig S4A). We next performed another WCB experiment with the different mutants and determined that the M25 strain binds human serum albumin (HSA), IgG, C4BP and α2-Macroglobulin (α2M) (Fig 4C). We found a decrease in the binding of both HSA and C4BP in the *Δemm* background. Although the major binder for IgG is the Mrp protein (23,27), we also observed a slight decrease in the triple KO compared to the *Δmrp* strain, suggesting that the M25 and/or Enn83 proteins also bind IgG. Finally, no difference was observed for α2M (Fig 4C), which was expected since it is bound by the GRAB protein (31). To further validate the interactions of the M25 protein with C4BP, HSA and IgG, we incubated a purified M25-His protein, immobilized on a Ni-NTA resin, with human serum. We found that all three proteins were eluted with the M25 protein (Fig 4D). Altogether, these data suggest that C4BP, HSA and IgG proteins could be involved in serum protection mediated by the M protein.

Finally, we determined their role in serum-mediated protection against φA25 infection. We used Intravenous Immunoglobulin (IVIg) preparation which contains mainly IgG and is used in treatment of invasive *S. pyogenes* infection (32,33). HSA transports fatty acids (FAs) and other molecules in the blood (34). Previous studies have demonstrated that the M1 protein is able to bind HSA which allows *S. pyogenes* to use HSA-bound fatty acids (23,35). The M25 wild-type strain was infected by the φA25 at a low MOI (0.001) in presence of C4BP at a concentration equivalent to 20% serum but we did not observe any significant protection (Fig S5A). We next used IVIg, native HSA (with fatty acids) but also fatty acid free HSA (HSA-FA) to mimic FAs depletion following uptake by *S. pyogenes*. They were tested either alone or in combination at a concentration equivalent to 20% serum on the M25 WT strain. We found a slight protection of HSA-FA against φA25 infection, while HSA favoured infection (Fig 5A). The addition of IVIg had a significant protective effect against φA25 infection which was enhanced in presence of HSA-FA (Fig 5A). By contrast, the IVIg-mediated protection was mitigated in presence of HSA (Fig 5A), suggesting that the content in FAs determines the effect of HSA. The effect of both HSA and HSA-FA was abolished in the *Δemm* background while the protective effect of IgG was maintained (Fig 5A). This is consistent with the interactions observed in the WCB assay where the M and Mrp proteins are the main binders of HSA and IgG, respectively (Fig 4C). We finally performed a WCB experiment using these proteins with the WT M25 strain and observed an increase in binding of HSA and HSA-FA in presence of IVIg (Fig 5B). Altogether, these data suggest that IgGs moderately participate in serum-mediated protection against φA25 infection in a Mrp protein-dependent manner while free fatty acid HSA (HSA-FA) participates in protection in a M-dependent manner.

**Fig 5.**
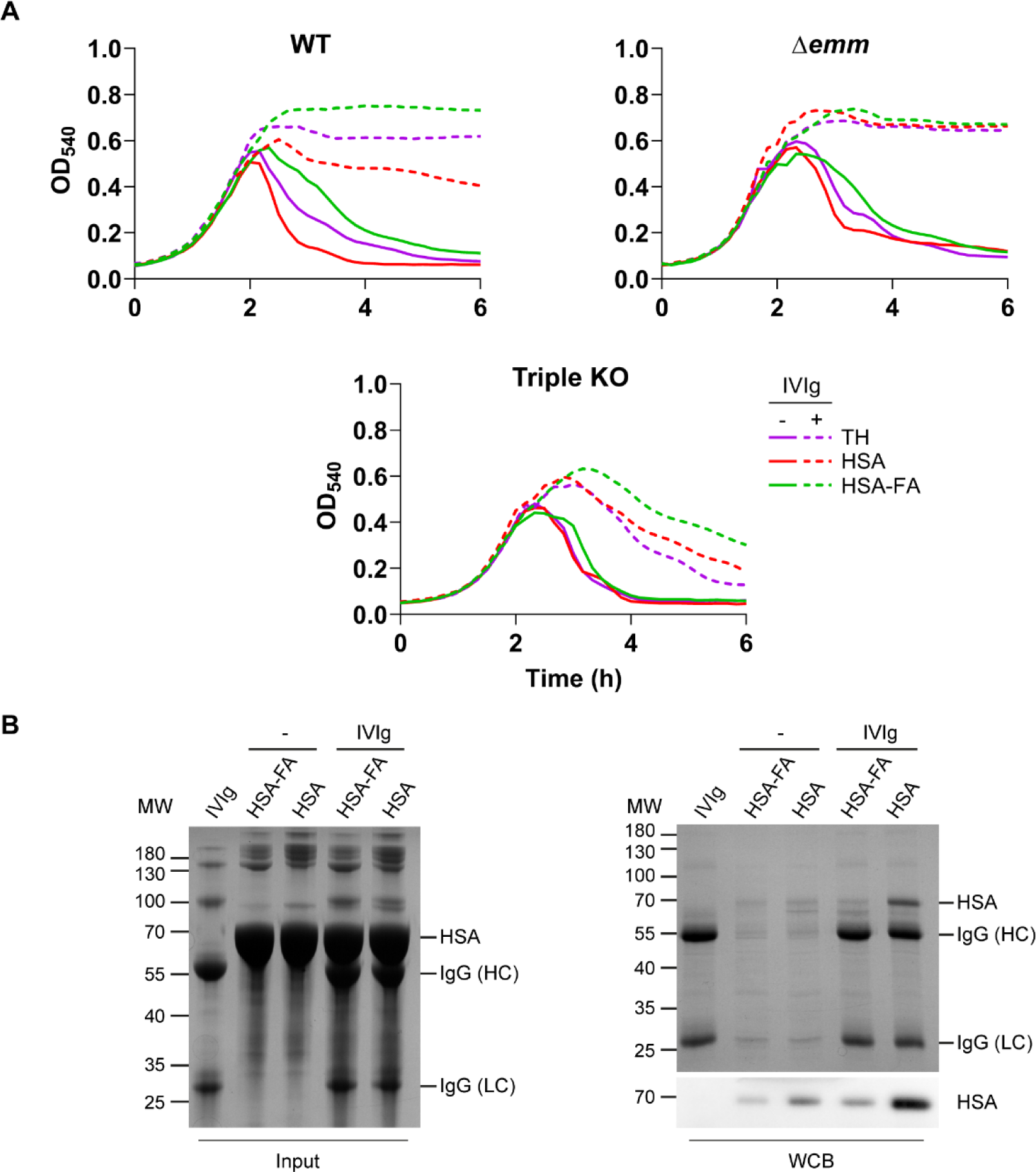
Role of HSA and IgG in protection against φA25 infection. **(A)** Collapse curves with φA25 (MOI=0.001) of *S. pyogenes* M25 WT, *Δemm* and triple KO mutant strains in TH medium alone (*purple*) or supplemented with 8 mg.mL^-1^ HSA (*red*) or HSA-FA (*green*) and containing (*solid lines*) or not (*dotted lines*) 3 mg.mL^-1^ IVIg. **(B)** SDS-PAGE/Coomassie blue staining profiles of IVIg, HSA, HSA-FA alone, or in combination, added (input, *left*) or bound (WCB, *right*) to the surface of the *S. pyogenes* M25 WT strain. SDS-PAGE/immunoblot analysis of the WCB using anti-HSA antibodies (*bottom right*). MW, *molecular weight* in kilodalton. Data in A are a representative of three independent experiments displayed in Fig S5B, including growth curves. Data in B are representative of three independent experiments.

## Discussion

The implementation of phage therapy in routine clinical practice requires safety and efficacy data. The phages selected for (pre)clinical trials are chosen based on *in vitro* results, without considering the environment in which they will be used. Growth rates, spatial and population structures, bystander microbiota or available resources are parameters that widely differ between the Petri dish and the infection site (8,36). Not surprisingly, the *in vitro* efficacy of several *Escherichia coli*, *Pseudomonas aeruginosa* and *Staphylococcus aureus* phages has failed to be transposed *in vivo* (37–40). Consequently, the success of phage therapy requires a better understanding of the phage-host interactions in *in vivo* and/or *ex vivo* conditions (8,41,42). In this work, we demonstrate that phage susceptibility of *Streptococcus pyogenes* is significantly altered in a human host-mimicking environment compared to conventional culture medium.

Phages-bacteria interactions in a relevant physiological context have recently begun to be characterized (39,40,43–48). For instance, both *E. coli* and *P. aeruginosa* increase the production of biofilms in mouse gut and human cystic fibrosis-like environment, respectively, reducing phages infection (39,46). *Vibrio cholerae* decreases the production of the phage ICP1 receptor (LPS O1-antigen) in response to intestinal bile acids and anaerobiosis, preventing phage predation (47). As well, recent works reported a negative effect of human plasma, particularly fibrinogen and synovial fluid, as well as the complement system, on *S. aureus* and *P. aeruginosa* phages infection (40,44,48). With our observations, all these data urge to isolate and select phages in conditions mimicking the “end-user” settings. Careful consideration must be given to the choice of phages, especially for ubiquitous bacteria residing in diverse ecological niches. Furthermore, the host-specificity of the pathogen is also an important criterion. In the case of *S. aureus*, diluted plasma from human and guinea pig were found to prevent phage infection (40). We found that mice and rabbit sera did not prevent φA25 infection, highlighting the difficulty to use *in vivo* models other than the NHPs or the human challenge model (25,49) to characterize phage efficacy against *S. pyogenes*. It also highlights the need for caution when interpreting the virulence data obtained in both mouse and rabbit models because some important interactions could be lost.

Besides the well-described genetic resistance to phage infection, like mutations of surface receptors or acquisition of defence systems (9), phenotypic resistance has been observed for different bacteria-phages pairs. It is a transient resistant state characterized by changes in bacterial genes expression interfering with phages adsorption (39,47,50,51). Notably, Lourenço and co-workers (39) highlighted that the mouse gut environment decreases expression of phages receptors in *E. coli*. Phase variation of phage receptors can also be used as a transient mechanism of resistance to phage infection, like described for *Bacteroidetes thetaiotaomicron* (52). Given that phenotypic resistance is a reversible and transient mechanism, it results in a lower fitness cost compared to genetic resistance, which is less readily reversible and long-lasting (53). We found that the addition of human serum to the M25 strain prevented φA25 adsorption within seconds, which excludes an effect on the expression of φA25 receptor(s). Moreover, after serum pre-incubation and washing, the protection was transient, and the phage ultimately collapsed the bacterial population. This is likely a consequence of the decrease in serum decoration of bacteria following their division. With the data obtained with plasma and fibrinogen for *S. aureus* (40), these are the first examples of a phenotypic resistance mechanism employing exogenous proteins rather than an adaptation of the transcriptome.

*S. pyogenes* is covered by a thick peptidoglycan which is decorated with group A carbohydrates, teichoic acid and LPXTG-motif proteins. The peptidoglycan is likely the receptor of φA25 (54). Therefore, any extensive coating of the peptidoglycan layer would prevent phage adsorption. Interestingly, two studies demonstrated that overexpressing the M protein or its removal by trypsin respectively increases and decreases resistance to infection of *S. pyogenes* by phages (including φA25 and its relatives) (55,56). The M25 strain carries the trio of M (*emm*) and M-like (*enn*, *mrp*) proteins coding genes (our unpublished data). Their deletion had no effect either positive or negative on the infection by the φA25 in absence of serum, suggesting that they did not take part in the adsorption. By contrast, in presence of serum, the deletion of the trio and the *emm* gene alone resulted in a notable increase in the collapse of the bacterial population. We demonstrated that the pre-incubation of serum with the wild-type (WT) bacteria depleted components required for protection. Conversely, the deletion of the *emm* gene partially restored protection conferred by the spent serum, likely because protective serum proteins were less depleted. Altogether, these results suggest that steric hindrance provided by an increase in protein composition (*i.e.* by co-opting serum proteins) confers resistance to *S. pyogenes* against φA25 infection. Yet, the composition of the serum proteins bound to the surface is important. Indeed, we found that heat-treatment of serum at 70°C changes the proportion of the different serum proteins which bind the bacterial surface and completely abolished the serum-mediated protection. Further works are needed to identify the serum proteins bound at the bacterial surface and *S. pyogenes* proteins involved in serum-mediated protection against φA25 infection.

The M and M-like proteins are known to interact with several human serum proteins. *In silico* search for binding motifs identified C4BP, IgG and HSA as potential partners of M25 in serum-mediated protection. Using whole cell binding (WCB) assay, we found that the *Δemm* strain is still able to bind IgG at its surface, while C4BP and HSA binding dropped significantly. We were further able to copurify these 3 serum proteins using GST-M25-His fusion. By infecting the wild-type strain in presence of these proteins, we observed that the C4BP did not affect φA25 infection. By contrast, using purified IgG (IVIg) and albumin without fatty acids (HSA-FA), we found a significant protection against phage infection, while HSA with fatty acids (HSA) had an opposite effect. We further demonstrated a synergistic binding of albumin and IgG on the surface of the M25 strain which correlates well with their effect during φA25 infection, *i.e.* additive (HSA-FA) or antagonistic (HSA) effect to IgG-mediated protection. The main effect of IgG was M protein-independent and likely due to Mrp which was found to be the main binder of IgG in the WCB assay. Since the purified M25 protein was found to interact with IgG, this synergistic binding is likely to occur by means of the M protein.

Our results demonstrate that the effects conferred by HSA and HSA-FA are M protein-dependent, since they were abolished in the *Δemm* background. The two opposite effects of HSA and HSA-FA were unexpected. The purified HSA used in this study is selectively precipitated from human plasma and further charcoal-treated to remove fatty acids (HSA-FA) (57,58). Hence, the phenotype observed with HSA-FA is unlikely to occur at the onset of φA25 infection in presence of serum. The *S. pyogenes* M1 strain binds HSA in a M protein-dependent manner which results in HSA-bound fatty acids uptake and a decrease of endogenous fatty acid synthesis (FASII pathway) (23,59). This process likely occurs in the M25 strain since we observed a downregulation of the FASII pathway in presence of HSA (our unpublished data). Consequently, the local production of HSA-FA at the bacterial surface is plausible as well as an effect on phage propagation or phage superinfection (*i.e.,* secondary infection of an infected bacterium). The determination of the crystal structure of both HSA forms highlighted distinct conformations with the rotation of two domains (60). These conformational changes might alter the binding affinity to the M protein and/or to other surface proteins. Understanding the fate of these proteins at the bacterial surface and therefore their effect on phage infection dynamics requires further experiments.

In conclusion, we found that human serum proteins protect *S. pyogenes* against infection by the phage φA25. This phage has an ancient lysogenic lifestyle (11,12,61). Since the virulence of *S. pyogenes* is tightly coupled to its prophages content (10,16), studying phages dissemination in presence of serum and HSA will help to understand its evolutionary journey. Like other human pathogens, *S. pyogenes* is adapted to different niches, either outside or inside the human host, like the skin, throat, blood, and deep tissues. All these niches are different from laboratory conventional medium. Knowing whether phages can efficiently kill their hosts in such diverse settings is key to the success of phage therapy.

## Material and methods

### Bacteria strains, phages and culture conditions

*Streptococcus pyogenes* M25 strain NCTC8306 (ATCC® 12204™) and phage φA25 (ATCC® 12204-B1™) were used in this study. Top10 *E. coli* strain (Invitrogen) was used for cloning and routinely grown at 37°C (30°C for protein production) with shaking at 220rpm in LB medium. *S. pyogenes* strains were plated on either blood agar plates or Todd-Hewitt supplemented with 0.5% yeast extracts (THY) plates. Bacteria were grown statically at 30°C (overnight growth) or at 37°C (day growth) in 5% CO_2_ atmosphere in autoclaved THY or in filtered TH medium. Growth curves with or without phages were determined in either a GENESYS™ 20 spectrophotometer (Thermofisher) in 1-mL cuvettes at 600nm or in a SpectroStar Nano plate reader (BMG LABTECH) at 540nm in sterile 96-well CELLSTAR® plates sealed with a sterile transparent film. Antibiotics used: ampicillin (100µg.mL^-1^ for *E. coli*); kanamycin (50µg.mL^-1^ for *E. coli* and 300µg.mL^-1^ for *S. pyogenes*); spectinomycin (100µg.mL^-1^ for both).

The φA25 was amplified using the M25 strain as described by P. Boulanger (62). Overnight culture was diluted to an optical density at 600nm (OD_600_) of 0.06 in autoclaved THY and exponentially grown (OD_600_≈0.4, which corresponds to 4x10^8^ CFUs/ml). The culture was supplemented with 1mM CaCl_2_/MgCl_2_ (THY^CM^) and infected for at least 2h with phages at a multiplicity of infection (MOI) between 0.1 and 1. Then, the bacteria were diluted in 4 volumes of THY^CM^ and further incubated overnight to allow complete lysis of the population. NaCl was dissolved in the lysed culture (500mM final conc.) and following a 1h incubation at room temperature (RT), bacterial debris and cells were removed by centrifugation at 8 000 g for 10 min at 4°C. Following overnight incubation at 4°C with 10% PEG 6000, the phages were centrifuged at 10 000 g for 15min at 4°C and resuspended in filtered TH^CM^. Phage A25 titer was determined by mixing (1:1) serial dilution of phages with 1/10^th^ of an overnight culture of M25 strain in THY^CM^. Spots were plated on THY^CM^ agar plates and incubated overnight (37°C + 5% CO_2_) to count plaque forming units (PFUs).

### Reagents

The following chemicals were used in this study: Mouse serum was purchased from Histoprime. Rabbit serum, Human Serum Albumin with or without fatty acids were purchased from Merck. IVIg (Multigam® 5%), a pool of purified IgGs, was purchased from Biotest Pharma GmbH. C4BP was purchased from Athens Research & Technology.

### Plasmid construction and mutant generation

All primers and plasmids used in this study are listed in Table S1-2, respectively. All generated plasmids were cloned using classical cloning or Goldengate assembly and validated by DNA sequencing. For both cloning, the different fragments were amplified with the PrimeSTAR® HS polymerase (Takara Bio) using specific primers flanked by restriction sites. For Goldengate cloning, *Bsa*I type IIS restriction site with 4-nt overhangs defining the order of assembly were used. The genes coding for the M25, Enn152 and Enn314 were cloned in the pGEX4T1 vector, with a 3’ his-tag and without their 5’ signal sequence and 3’ LPXTG motif, and then transformed in *E. coli* Top10 for expression. Deletion mutants were generated using a suicide vector for *S. pyogenes* consisting in the non-replicative pUC18 vector expressing spectinomycin (Schiavolin *et al.*, in preparation) and containing 1-kb flanking regions of the gene to deleted which surround the *aphA3* gene (kanamycin resistance). After PCR purification and assembly reaction, the products were transformed in electrocompetent *E. coli*. Transformants were selected on spectinomycin LB agar plates. Suicide plasmids (500-1000 ng) were transformed in electrocompetent *S. pyogenes* using the sucrose method (63). Transformants were selected on THY agar plates containing kanamycin. Clones were then isolated on either spectinomycin or kanamycin THY agar plate and kan^R^ spec^S^ clones (double allelic exchange) were screened and validated by PCR.

### Phage experiments

All phage infection experiments were performed in presence of 1mM CaCl_2_/MgCl_2_ (CM). Phage adsorption was performed as described by A. Kropinski (64). The M25 strain was exponentially grown (OD_600_ of 0.4 - 0.6) in TH medium. The serum was added at a final concentration of 20% just before infection. Bacteria and CTL medium were inoculated with phages at a MOI of 0.001 and incubated at 37°C in Lab Armor™ beads bath. Samples were collected every minute, diluted in ice-cold THY supplemented with CHCl_3_ and vortexed for 10sec. Samples were finally mixed with an equal volume of a 1/10^th^ dilution of an M25 overnight and spotted on THY^CM^ agar plates for PFU counting.

For CFU/mL and MOI determination, bacteria were grown in THY medium to exponential, early and late stationary phases. OD_600_ was measured and bacteria diluted for counting, before and after sonication at 4°C for 15min in a Bioruptor® (Diagenode), to determine Chain (ChFU) and Colony (CFU) Forming units per mL at an OD_600_=1, respectively. For φA25 infection, bacteria were exponentially grown (mid-log phase). Following sonication, they were infected at 37°C with different MOIs. The adsorption was stopped after 10min by serial dilution in ice-cold THY and bacteria were spotted on THY^CM^ agar for CFU counting.

One-step growth curves were determined as described by Chevallereau and colleagues (65). The M25 strain was exponentially grown (OD_600_ of 0.4 - 0.6) in TH medium. Bacteria were infected 8 minutes at a MOI of 0.01 at 37°C and diluted 100 times in TH at 37°C. Samples were collected at different time points, diluted 20 times in ice-cold THY supplemented or not with CHCl_3_ and vortexed for 10sec. Samples were finally mixed with the M25 strain for PFU counting. The eclipse (+CHCl_3_, free phages) and latency (-CHCl_3_, free phages + infective centres) periods were then calculated and correspond to the time needed to detect 5% of total PFUs for each condition. The burst size was determined as follow: [PFU _(+CHCl3),t=xmin_]/ ( [PFU _(-CHCl3),t=10min_]-[PFU _(+CHCl3),t=10min_]).

Phage infection was performed at 37°C either in a beads bath with OD_600_ measurement using a Genesys 20 spectrophotometer (Thermo scientific) or in 96-well plates using a SPECTROstar® Nano (BMG Labtech) with OD_540_ measurement every 10 minutes. Two different MOIs were used, except where otherwise stated. Phages were added at (i) low MOI (0.001) to bacteria diluted from overnight culture at an OD_600_ of 0.06 to quantify the propagation of the phages in the bacterial population; (ii) high MOI (5 to 25) to bacteria previously exponentially grown to infect the whole bacterial population and perform subsequent analysis.

For pre-incubation experiments with human serum, the M25 strain was exponentially grown (OD_600_ of 0.4 - 0.6) in TH medium. Bacteria were then divided into two aliquots, pelleted at 5000×g for 10 minutes, and resuspended in either 2mL TH medium or 100% human serum. After 30 minutes of incubation at 37°C, the bacteria were washed once with TH medium, OD_600_ was adjusted at 0.4 and bacteria were mixed with phages at different MOIs. Depleted serum was further filtered and used at 10% final concentration with bacteria at an OD_600_ of 0.06 and phages at low MOI. Serum was also depleted with the WT and *Δemm* strains in a second set of experiments.

### Whole cell binding assay

The M25 and its mutant strains were exponentially grown (OD_600_ of 0.4 - 0.6) in THY medium, pelleted at 5000×g for 10 minutes and washed in PBS. Bacterial pellets (10^10^ CFU) were resuspended in 500 µL of human serum (heat-treated or not at 70°C) and incubated with gentle agitation for 2 hours at 25°C. Bacteria were pelleted and washed three times with ice-cold PBS supplemented with Tween20 0.05%. The pellets were resuspended with 50 µL of recommended iD PAGE sample buffer 1X, centrifuged at 9000×g for 2 minutes and the supernatants were kept for further use at -20°C.

### Copurification experiments

*E. coli* Top10 strains expressing the pGEX4T1-his constructs were exponentially grown (OD_600_ of 0.6 - 0.8) in LB medium. IPTG was added at 1 mM and bacteria were further incubated at 30°C for 3 hours. Bacteria were pelleted at 4000×g for 15 minutes at 4°C. Pellets (from 200mL) were vortexed in 4 mL of ice-cold equilibration/wash (EW) buffer (50 mM sodium phosphate, 300 mM sodium chloride, 10 mM imidazole; pH 7.4) supplemented with lysozyme at 0.2 mg.mL^-1^. Bacteria were then lysed with 1 g of lysing matrix B using a FastPrep-24™ 5G (MP Biomedicals). Lysates were centrifuged at 8000×g for 30 minutes at 4°C and supernatants were kept on ice for further use. For each lysate, 650 µL of HisPur^TM^ Cobalt resin (Thermofisher) was centrifuged at 700×g and washed twice with 1 mL of ice-cold EW buffer. Resins and lysates were then mixed and incubated with gentle agitation for 1 hour at 25°C. After three washes with 5 mL of ice-cold EW buffer, each resin was incubated for 2 hours at 25°C with 20U thrombin to remove the N-terminal GST tag. Resins were further washed three times with EW buffer and incubated with 5 mL of a 1/10^th^ dilution of human serum in PBS and incubated with gentle agitation overnight at 4°C. Finally, resins were washed 10 times with 5 mL ice-cold EW buffer. Proteins were then eluted with ca. 500µl of sample buffer 1X with gentle agitation for 10 minutes at 25°C and kept at -20°C for further analysis.

### SDS-PAGE and western blotting

SDS-PAGE were run on iD PAGE gel 10% (Eurogentec) with MOPS buffer and bands detected with Coomassie blue staining. The western blot analysis was performed on PVDF membrane using rabbit anti-IgG-HRP (Dako), mouse anti-Albumin (Tebu-bio), rabbit anti-C4BPA (Merck), or rabbit anti-A2M (Invitrogen) primary antibodies and anti-mouse-HRP (Merck) and anti-rabbit-HRP (Tebu-bio) secondary antibodies when required. Immunoreactive bands were detected by chemiluminescence using the Clarity™ Western ECL Substrate (Bio-Rad) and an Amersham Imager 600 (GE Healthcare Life Sciences).

### Human samples and Ethics statement

Human venous blood was obtained from healthy volunteers in dry tubes. After clotting and centrifugation, sera were pooled, aliquoted and freeze-stored before use.

Volunteers were recruited by a passive advertising campaign within the Faculty of Medicine at the Université Libre de Bruxelles. Written consent was obtained from each volunteer before blood collection. Blood samples were anonymized, and no donor information were retained. The protocol has been validated by the ethics committee of the H.U.B Erasmus Hospital of Brussels (ref: P2015/398).

### Statistics

All statistical tests were performed using GraphPad Prism 10. Before performing two-tail unpaired t test and ANOVAs, we determined the normal distribution of the data using a Shapiro-Wilk test. Plots were generated using GraphPad Prism 10.

## Acknowledgments

We thank members of the Microbiology Laboratory and Pr. Anne Op De Beeck for helpful discussions. The work was supported by a Belgian Fonds National de la Recherche Scientifique (FNRS, http://www.fnrs.be) research grant (PDR T.0255.16). L.S. was founded by a Post-Doc fellowship from FNRS. J.S. was founded by a PhD fellowship from FRIA (FNRS).

## Supplementary figures

**Fig S1, related to Fig 1.**
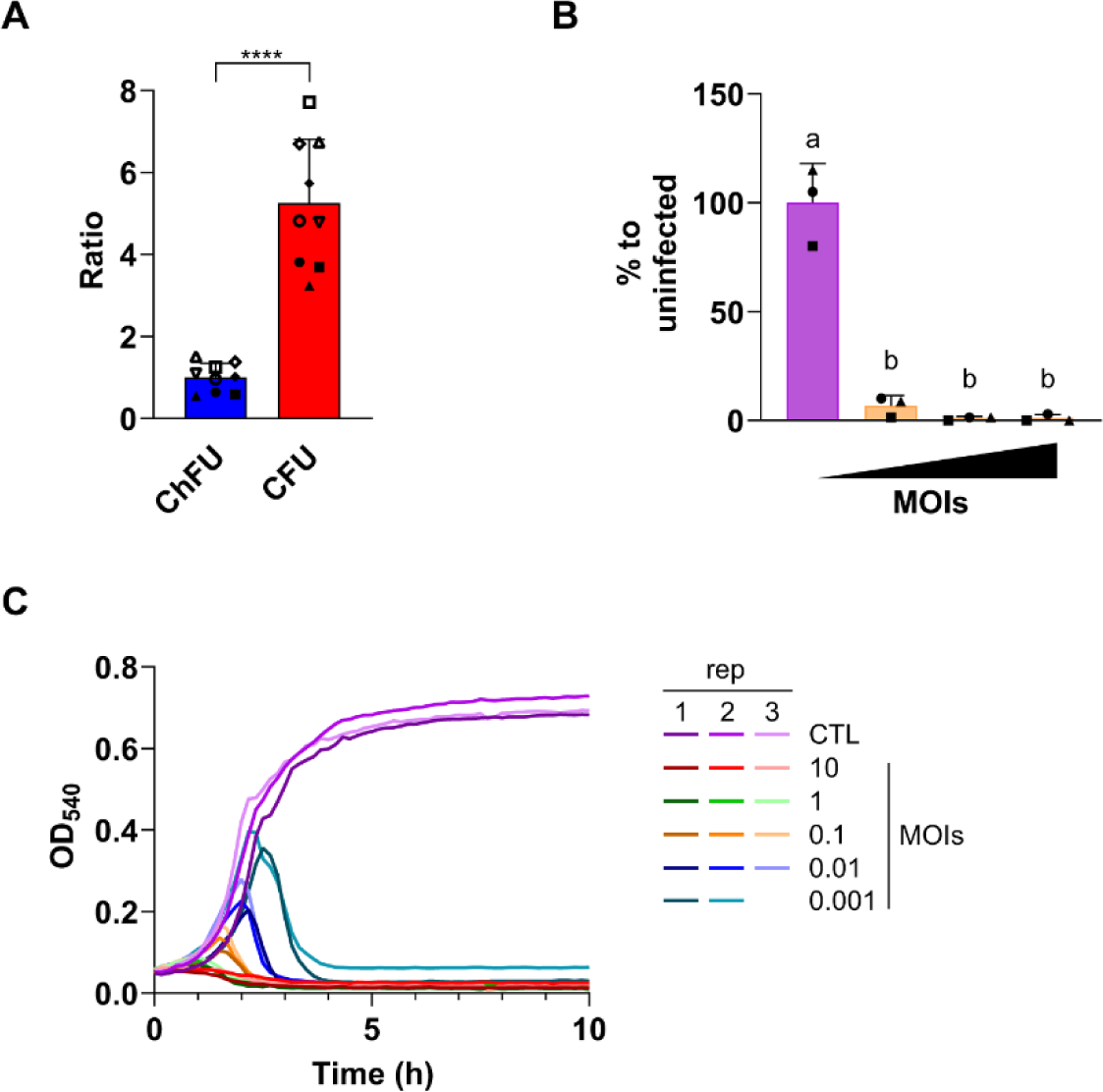
Phage φA25 – M25 strain infection parameters. **(A)** Number of *S. pyogenes* M25 WT bacteria per chain after sonication. Each value is expressed as a ratio to the mean ChFU.mL^-1^ at an OD_600_=1 (n=9). **(B)** Number of φA25 (MOI) required to eradicate the sonicated *S. pyogenes* M25 population. Values are expressed as a percentage to the mean CFU of uninfected bacteria (MOI=0) (n=3). **(C)** Replicates of curves displayed in Fig 1D with a gradient colour and the same colour scheme. Data (A-B) are displayed as mean ± SD of at least three independent experiments. Statistical tests. Unpaired t test (A): ****, p<0.0001 (t=10.16, df=8); One-way ANOVA (B): a-b, p<0.0001 (t=81.40, df=3).

**Fig S2, related to Fig 2.**
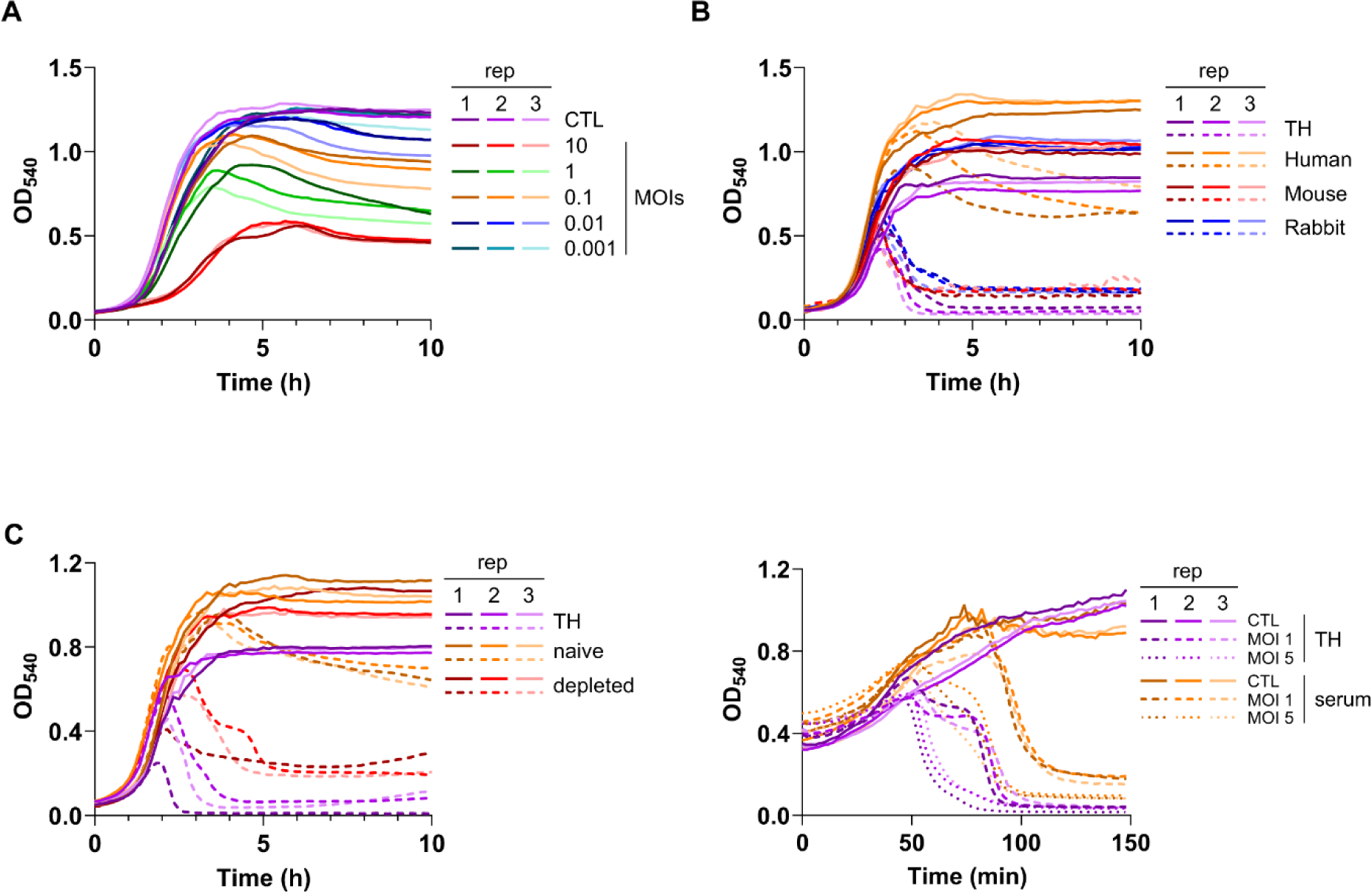
Human Serum protects against φA25 infection. **(A-C)** related to Fig 2A-C, respectively, with a gradient colour and the same colour scheme. Data are displayed as individual replicates of three independent experiments.

**Fig S3, related to Fig 3.**
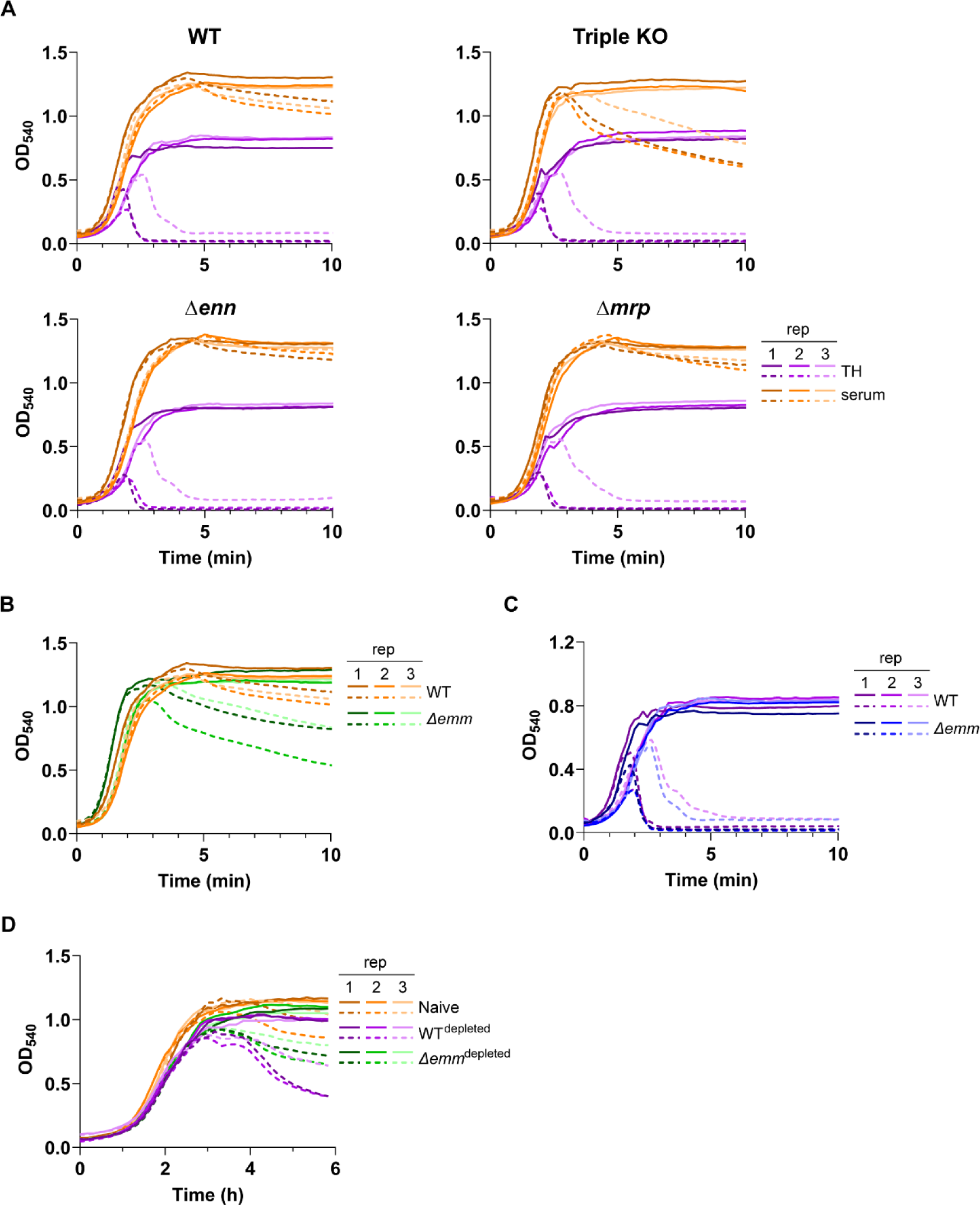
The M protein contributes to serum-mediated protection against φA25 infection. **(A)** Growth (*solid line*) and collapse (*dotted line*) curves with φA25 (MOI=0.01) of *S. pyogenes* M25 WT and mutant (*Δenn*, *Δmrp* and triple KO) strains in TH medium alone (*purple*) or supplemented with 20% human serum (*orange*). **(B and D)** related to Fig 3A-B, respectively, with a gradient colour and the same colour scheme. **(C)** Growth (*solid line*) and collapse (*dotted line*) curves with φA25 (MOI=0.01) of *S. pyogenes* M25 WT (*purple*) and *Δemm* (*blue*) strains in TH medium, related to Fig 3A. Data are displayed as individual replicates of three independent experiments.

**Fig S4, related to Fig 4.**
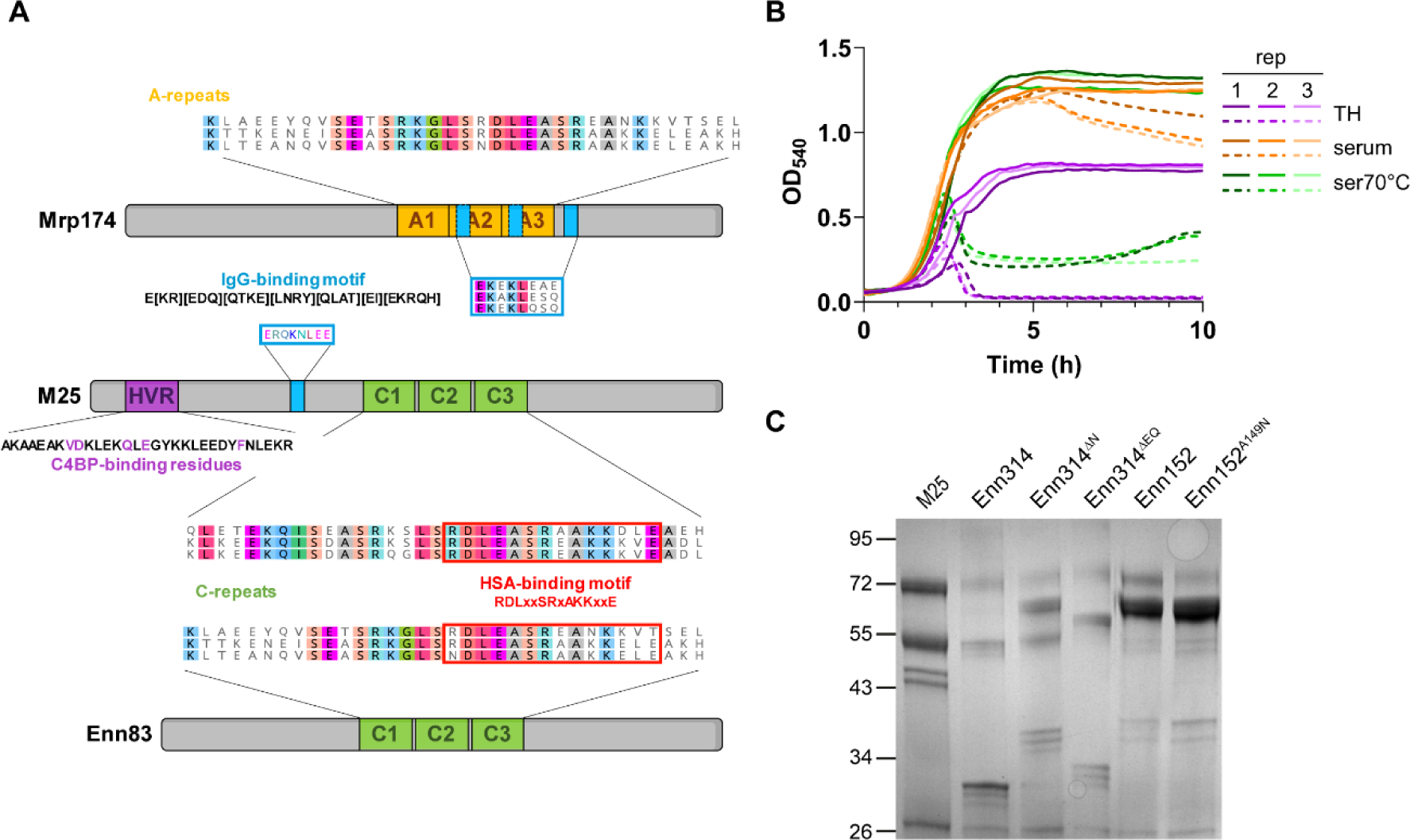
Role of serum proteins in protection against φA25 infection. **(A)** Schematic representation of the three M and M-like proteins of *S. pyogenes* M25 strain with the different binding motifs. **(B)** related to Fig 4A with a gradient colour and the same colour scheme. Data are displayed as individual replicates of three independent experiments. **(C)** SDS-PAGE/Coomassie blue staining of the elution of purified proteins as indicated, related to Fig 4D.

**Fig S5, related to Fig 5.**
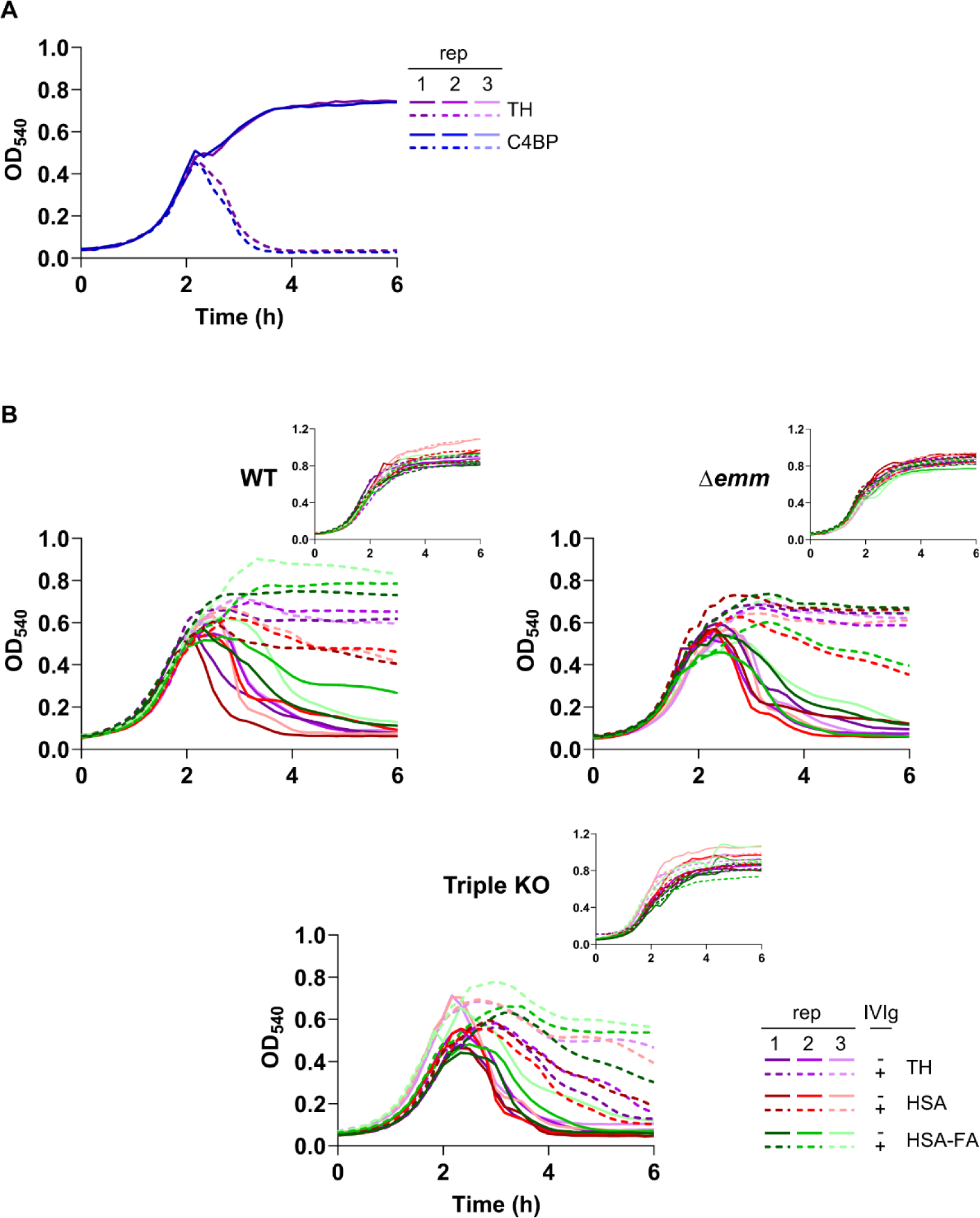
Role of HSA and IgG in protection against φA25 infection. **(A)** Growth (*solid line*) and collapse (*dotted line*) curves with φA25 (MOI=0.001) of *S. pyogenes* M25 WT strain in TH medium alone (*purple*) or supplemented with 40 µg.mL^-1^ C4BP (*blue*) (n=1). **(B)** Growth (*inset*) and collapse curves related to Fig 5A with a gradient colour and the same colour scheme. Data are displayed as individual replicates of three independent experiments.

## Notes

### Competing Interest Statement

The authors have declared no competing interest.

